# R2R3-MYB EVER links emission of volatiles with epicuticular wax biosynthesis in petunia petal epidermis

**DOI:** 10.1101/2023.06.15.545203

**Authors:** Oded Skaliter, Dominika Bednarczyk, Ekaterina Shor, Elena Shklarman, Ekaterina Manasherova, Javiera Aravena-Calvo, Shane Kerzner, Alon Cna’ani, Weronika Jasinska, Tania Masci, Gony Dvir, Orit Edelbaum, Ben Spitzer-Rimon, Yariv Brotman, Hagai Cohen, Alexander Vainstein

## Abstract

The epidermal cells of petunia flowers are the main site of volatile emission. However, data on the mechanisms underlying the release of volatiles into the environment are lacking. Here, using cell-layer-specific transcriptomic analysis, reverse genetics by VIGS and CRISPR, and metabolomics we identified EPIDERMIS VOLATILE EMISSION REGULATOR (EVER)—a petal adaxial epidermis-specific MYB activator that affects the emission of volatiles. Using a three-step viral-based CRISPR/Cas9 editing system, *ever* knockout lines were generated and together with transient suppression assays, revealed EVER’s involvement in the repression of low-vapor-pressure volatiles. Internal pools and annotated scent-related genes involved in production and emission were not affected by EVER. RNA-Seq analyses of petals of *ever* knockout lines and *EVER*-overexpressing flowers revealed enrichment in wax-related biosynthesis genes. LC/GC-MS analyses of petal epicuticular waxes revealed substantial reductions in wax loads in *ever* petals, particularly of monomers of fatty acids and wax esters. These results implicate EVER in the emission of volatiles by fine-tuning the composition of petal epicuticular waxes. Thus, we reveal a petunia MYB regulator that interlinks epicuticular wax composition and volatile emission, thus unraveling a new regulatory layer in the scent-emission machinery in petunia flowers.

## Introduction

Plants developed the ability to synthesize phenylpropanoids ca. half a billion years ago, when they first started colonizing land, helping them adapt to the new environmental stresses accompanying terrestrial life (Weng and Chapple, 2010; Banks et al., 2011). During the course of evolution, the phenylpropanoid pathway has diversified through the emergence of new branches, allowing plants to synthesize a multitude of metabolites from the precursor phenylalanine, including volatiles (Weng et al., 2012). The latter play an essential role in plants’ life cycle by allowing them to interact with biotic and abiotic factors in their surrounding environment, for example, by attracting pollinators or as airborne warning signals. Some volatiles are aroma compounds that contribute to food flavor and may carry health benefits; these are highly valuable to the food, beverage and pharma industries (Skaliter et al., 2022). Volatile molecules are characterized by low molecular weight and high vapor pressure, enabling them to evaporate at ambient temperature. *Petunia* x *hybrida* is a common model plant for scent studies due to its large flowers, which produce and emit phenylalanine-derived volatiles (PDVs) in significant amounts. These include benzenoids (C6–C1 backbone), phenylpropanoid-related compounds (C6–C2) and phenylpropenes (C6–C3) (Muhlemann et al., 2014). Furthermore, petunia is amenable to reverse genetic techniques such as virus-induced gene silencing (VIGS) and gene editing by CRISPR/Cas9, and sequenced genomes of its ancestors *Petunia axillaris* and *Petunia inflata* are available (Spitzer et al., 2007; Bombarely et al., 2016; Chopy et al., 2020; Lin and Jones, 2022). In the last two decades, extensive research has led to the identification and characterization of most of the enzymes responsible for the synthesis of the aromatic amino acid (AAA) precursors of the pathway(Lynch and Dudareva, 2020) and of dedicated PDV-biosynthesis enzymes (Muhlemann et al., 2014; Huang et al., 2022; Skaliter et al., 2022). A number of transcription factors (TFs) regulating this pathway have been identified (Verdonk et al., 2005; Spitzer-Rimon et al., 2010; Colquhoun et al., 2011; Spitzer-Rimon et al., 2012). Among these, a triad of MYBs—ODORANT1 (ODO1), and EMISSION OF BENZENOIDS I and II (EOBI and EOBII)—have been shown to work in concert and coregulate each other, in addition to their direct regulation of AAAs and scent-related biosynthesis genes (Verdonk et al., 2005; Spitzer-Rimon et al., 2010, 2012; Boersma et al., 2022). Upstream of the MYB triad are two members of the GRAS family: PHENYLPROPANOID EMISSION-REGULATING SCARECROW-LIKE (PES), shown to activate *ODO1*’s promoter (Shor et al., 2023a), and Ph-DELLA—a suppressor of gibberellin (GA) signaling that can activate *EOBII* (Ravid et al., 2017). In addition to GA, auxin and ethylene have also been shown to be involved in the regulation of floral scent. Auxin, like GA, acts at bud stages, negatively affecting PDV production by altering plastid development and phenylalanine levels (Ravid et al., 2017; Lynch et al., 2020). Ethylene acts at late stages of flower development, partially through ETHYLENE RESPONSE FACTOR 6 (ERF6) which interacts with EOBI’s N terminus, thus hindering activation of its target genes (Underwood et al., 2005; Liu et al., 2017). Like ERF6, the MYBs Ph-MYB4 and EMISSION OF BENZENOIDS V (EOBV) negatively regulate scent. Ph-MYB4 affects carbon flux toward C6–C3 compounds by downregulating *cinnamate 4-hydroxylase* (Colquhoun et al., 2011). EOBV’s mode of action is still unknown (Spitzer-Rimon et al., 2012). Recently, UNIQUE PLANT PHENYLPROPANOID REGULATOR (UPPER), with a unique RING–KINASE–WD40 domain structure, was revealed to be involved in negative regulation of PDVs in petunia flowers, adding another layer to the complex network of scent regulation (Shor et al., 2023b).

Following their production in the cytosol, endoplasmic reticulum, peroxisome or mitochondria, PDVs may be compartmentalized in cellular organelles such as the vacuole if glycosylated (Cna’ani et al., 2017), or released to the environment or transported through the plasma membrane and cell wall of epidermal cells, and through the overlying cuticle (Adebesin et al., 2017; Liao et al., 2023). The cuticle is multilayered and built from the cutin polymer and waxes. Cutin is a mixture of very-long-chain C16 and C18 fatty acid derivatives with terminal and mid-chain hydroxy, epoxy and carboxy functional groups (Cohen et al., 2019; Philippe et al., 2020). Intracuticular waxes are dispersed within the cutin matrix, and epicuticular waxes are deposited on top of the cuticle (Arya et al., 2021; Sarkar et al., 2023). Waxes consist of very-long-chain fatty acids (VLCFAs, typically >C20), and their derivatives belonging to various biochemical classes, including primary and secondary alcohols, aldehydes, alkanes, ketones and esters (Samuels et al., 2008; Buschhaus and Jetter, 2011). The cuticle has been recently suggested to play a regulatory role in volatile production and emission via suppression of Ph*-ABCG’s*— transporters involved in the transport of cutin monomers (Liao et al., 2021). Of note, wax-biosynthesis genes, as well as those from the phenylpropanoid pathway that generate flavonoids and PDVs, are preferentially expressed in epidermal cells (Kolosova et al., 2001; Scalliet et al., 2006; Lee and Suh, 2015; Lee et al., 2016; Verweij et al., 2016; Negin et al., 2022). This specialization of the epidermis, namely high emission of volatiles and synthesis of cuticular waxes, places it as an ideal layer for studying the genes involved in these processes. Furthermore, as shown in petunia, the adaxial epidermis, relative to the abaxial one, has been shown to be the main site for the emission of PDVs (Skaliter et al., 2021). In this research, we characterized the transcriptomic profile of the adaxial epidermis of petunia cv. Mitchell. Screening for adaxial epidermis-enriched TFs yielded Peaxi162Scf00777g0052, which was named *EPIDERMIS VOLATILE EMISSION REGULATOR* (*EVER*). Transient downregulation using VIGS and viral-based CRISPR/Cas9-induced knockout of *EVER* led to increased emission of floral PDVs, revealing it as a negative regulator of this trait. RNA-Seq and LC/GC–MS analyses performed on *ever* knockout lines demonstrated that the transcriptional regulator EVER controls epicuticular wax buildup on the petal, revealing the importance of wax composition in floral scent emission.

## Results

### Transcriptomic profile of the adaxial petal epidermis vs. whole petal of petunia cv. Mitchell

To study the transcriptome profile of the adaxial epidermis, this layer was isolated from corollas of cv. Mitchell (W115) as reported previously (Skaliter et al., 2021) and used for RNA-Seq analysis. Petal tissues from flowers 1 day postanthesis (1DPA) were chosen for the analysis as scent production is high and transcripts of scent-related genes are enriched at this developmental stage (Shor et al., 2023b). As a reference, we extracted RNA from whole petal tissue (WPT), i.e., both epidermal layers and the mesophyll (Skaliter et al., 2021). Analysis of the transcriptome revealed 20,875 expressed genes. Using fold-change ≥1.5 and false discovery rate correction (FDR) <0.05 as a cutoff, we found 1393 (6.67%) genes that were more highly expressed in the adaxial epidermis than in the WPT (Fig. 1a, Supplemental Data S1 and S2). As expected, among these genes were *PH3* (Peaxi162Scf00472g00077), *PH5* (Peaxi162Scf00177g00620) and *PH4* (Peaxi162Scf00349g00057), exhibiting a 1.6-, 1.7- and 2.5-fold higher expression, respectively (Quattrocchio et al., 2006; Verweij et al., 2008, 2016). Two genes with higher expression in the adaxial epidermis (Peaxi162Scf00069g02239/g02248), annotated as “Epidermis-specific secreted glycoprotein EP1”, were shown by *in situ* hybridization in carrot (*Daucus carota*) to be specific to the epidermal layer (Van Engelen et al., 1993). Furthermore, homologs of Arabidopsis genes known to be exclusively expressed in the epidermis, such as *LONG-CHAIN ACYL-COA SYNTHETASE* (*LACS*) (Peaxi162Scf00786g00434) and *PROTODERMAL FACTOR 2* (*PDF2*) (Peaxi162Scf00079g00092), also showed higher expression in the adaxial epidermis (Suh et al., 2005; Kamata et al., 2013).

**Figure 1.**
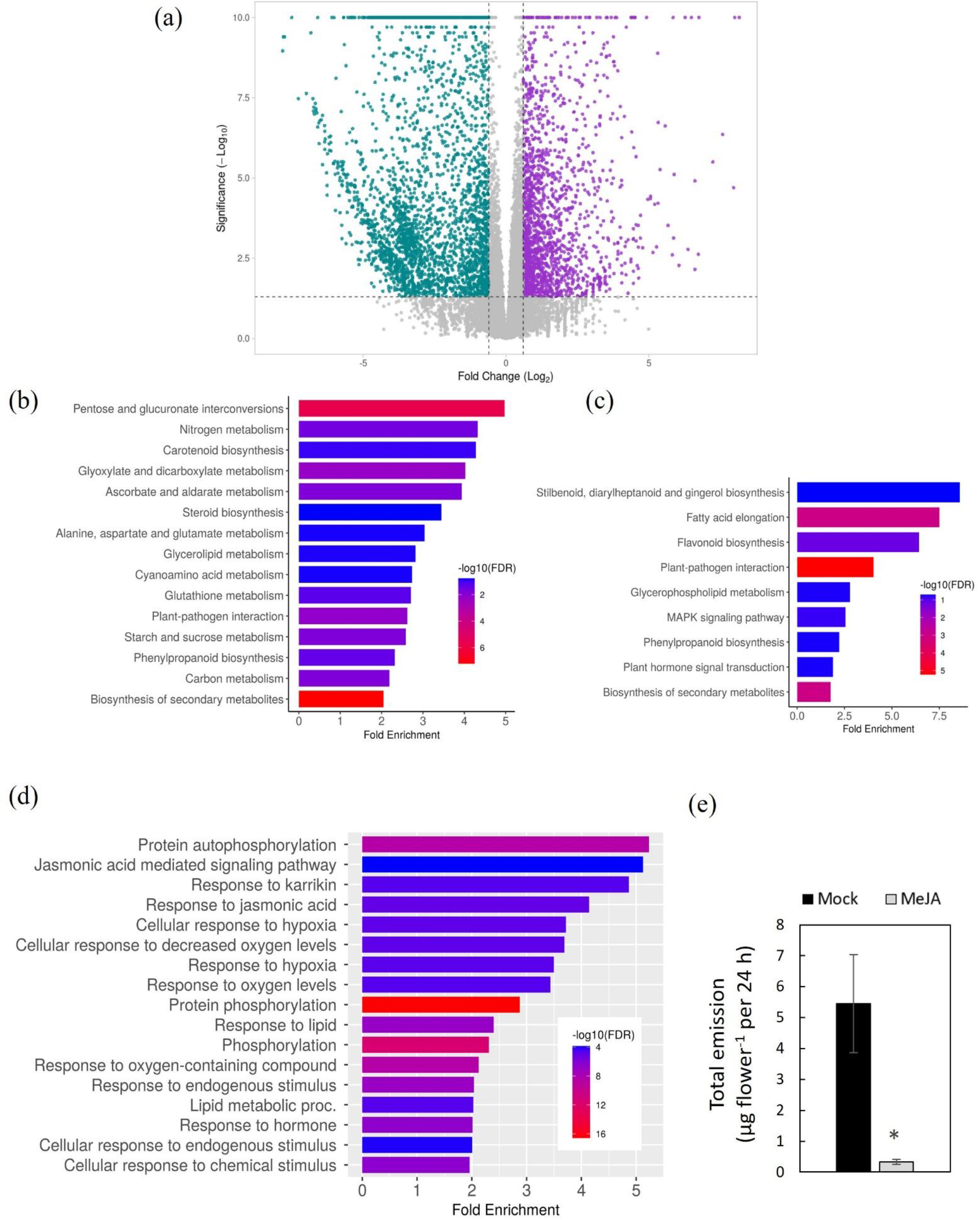
Transcriptomic analysis of the adaxial epidermis vs. whole petal tissue (WPT) of petunia cv. Mitchell. (a) Volcano plot for differential gene expression. Scattered dots represent genes. Differentially expressed genes with higher expression in the adaxial epidermis are presented in purple, those with higher expression in the WPT are in cyan, and those that are not differentially expressed are in gray. (b, c) KEGG enrichment analysis of differentially expressed transcripts that are more highly expressed in WPT (b) or in adaxial epidermis (c). (d) Enriched GO “Biological process” terms in the adaxial epidermis. (e) Effect of methyl jasmonate (MeJA) application on emission of volatiles from petunia flowers. Following 36 h treatment (with or without MeJA), flowers were analyzed by dynamic headspace conducted for 24 h, followed by GC-MS analyses. Data are means ± SEM (n = 5). Significance of differences (\**P* ≤ 0.05) between treatments was calculated by Student’s t-test.

In the WPT, 1771 (8.48%) genes showed higher expression than in the adaxial epidermis (Fig. 1a, Supplemental Data S1 and S3). These included mesophyll-specific genes such as the homolog of *FBase* from wheat (Peaxi162Scf01302g00031) (Sade et al., 2014), with 16-fold higher expression in the WPT. Moreover, 17 genes with higher expression in the WPT were annotated as photosynthesis-related, for example: *VIOLAXANTHIN DE-EPOXIDASE* (Peaxi162Scf01011g00215), *PHOTOSYSTEM II 22 KDA PROTEIN* (Peaxi162Scf00051g00910) and *CHLOROPHYLL A/B BINDING PROTEIN* (*CAB*) (Peaxi162Scf00047g02128), in agreement with the photosynthetic activity of mesophyll parenchyma cells in petunia petals (Weiss et al., 1988; Vainstein and Sharon, 1993; Cavallini-Speisser et al., 2021). Quantitative real-time PCR (qPCR) analyses were performed on *ACTIN* (Peaxi162Scf00023g02024, not differentially expressed), *CHALCONE SYNTHASE a* (*CHS*) (Peaxi162Scf00047g01225, not differentially expressed, from the flavonoid pathway), *CAB* (higher expression in WPT) and *PH4* (higher expression in adaxial epidermis) to confirm the authenticity of the transcriptome (Supplemental Fig. S1).

Scent-related genes from the phenylpropanoid pathway that showed higher expression in the adaxial epidermis than in the WPT included *CINNAMATE 4*-*MONOOXYGENASE b* (*C4H*, Peaxi162Scf00390g00225), *CONIFERYL ALCOHOL ACETYLTRANSFERASE* (*CFAT*, Peaxi162Scf00474g00217), *4-COUMARATE:COA LIGASE* (Peaxi162Scf00314g00086), *EOBV* (Peaxi162Scf00362g00831), Ph*-ABCG11a* (Peaxi162Scf01390g00033), Ph*-ABCG12b* (Peaxi162Scf00004g03212), *PHENYLALANINE AMMONIA-LYASE 1* (*PAL1*, Peaxi162Scf00858g00215) and *PAL2* (Peaxi162Scf00123g00096). AAA-biosynthesis genes that showed higher expression in the adaxial epidermis than in the WPT were *AROGENATE DEHYDRATASE 4* (Peaxi162Scf00147g00613), *CHORISMATE MUTASE 1* (*CM 1,*Peaxi162Scf00166g00931) and *3-DEOXY-D-ARABINO-HEPTULOSONATE-7-PHOSPHATE 1 SYNTHASE 1* (*DAHPS 1,* Peaxi162Scf00030g01715). In addition, a putative *CINNAMYL ALCOHOL DEHYDROGENASE* (Peaxi162Scf00623g00411) was upregulated in the adaxial epidermis as well. The scent regulators *ODO1* (Peaxi162Scf00002g00037), *EOBII* (Peaxi162Scf00080g00064), *EOBI* (Peaxi162Scf00129g01231), and Ph*-MYB4* did not show higher expression in the adaxial epidermis vs. the WPT.

To further characterize the RNA-Seq data, Kyoto Encyclopedia of Genes and Genomes (KEGG) pathway-enrichment analysis (FDR <0.2) was performed. Supporting the mesophyll’s role in petal nutrition and metabolism (Cavallini-Speisser et al., 2021), “Pentose and glucuronate interconversions”, “Carbon metabolism”, “Nitrogen metabolism” and “Starch and sucrose metabolism” were enriched in the WPT (Fig. 1b). Furthermore, “Carotenoid biosynthesis” and “Steroid biosynthesis” pathways, which can be attributed to photosynthetic activity of the mesophyll (Weiss et al., 1988; Vainstein and Sharon, 1993), were also enriched. “Flavonoid biosynthesis” and “Stilbenoid diarylheptanoid and gingerol biosynthesis”, as opposed to the general “Phenylpropanoid biosynthesis” pathways were enriched only in the adaxial epidermis (Fig. 1, b and c). As expected, “Fatty acid elongation” was also enriched in the epidermis, which can be explained by formation of the cuticle in this tissue (Suh et al., 2005). To obtain more information about the transcriptomic profile of the genes with higher expression in the adaxial epidermis than in the WPT, we also performed biological process gene ontology (GO) enrichment (FDR <0.05). Among the enriched processes was “Lipid metabolic process”, which encompasses wax and cuticle biosynthesis. In addition, “Response to hormone” and specifically “Response to jasmonic acid (JA)” were also enriched (Fig. 1d). None of these processes were enriched in GO for the WPT (Supplemental Fig. S2a). The biological process “Response to hormone” was enriched in the adaxial epidermis with 83 genes. Among them were genes related to phytohormone production/mode of action, shown to be involved in PDV biosynthesis in petunia flowers: auxin, ethylene and GA (Underwood et al., 2005; Ravid et al., 2017; Lynch et al., 2020; Patrick et al., 2021).For example, genes related to GA biosynthesis and signal transduction included homologs of *GIBBERELLIC ACID INSENSITIVE* (*GAI*; Peaxi162Scf00361g00101) (Sun, 2011) and the GA catabolic gene *GA2OX2* from Arabidopsis (Peaxi162Scf00111g00923) (Lantzouni et al., 2017). Interestingly, JA was not directly linked to scent production in petunia. Among the 20 genes involved in the enriched JA processes were: a putative *JASMONATE-INDUCED OXYGENASE 1*, involved in JA catabolism (Peaxi162Scf00014g00041) and several JASMONATE-ZIM-DOMAIN PROTEINs: JAZ-like *JAZ4* (Peaxi162Scf00314g00330) and *JAZ7* (Peaxi162Scf00317g00117), whose expression has been shown to increase toward anthesis in petunia (Tian et al., 2019). Conversely, a homolog of *CORONATINE INSENSITIVE 1* (Peaxi162Scf00401g00117), a component of the JA receptor complex, was strongly downregulated in the adaxial epidermis as compared to the WPT. To determine whether JA is involved in the regulation of PDV biosynthesis in petunia, flowers were treated with methyl jasmonate (MeJA), the volatile form of JA. Dynamic headspace and GC–MS analyses of MeJA-treated flowers, as compared to control mock treatment, led to a strong decrease in total emission levels (ca. 16-fold), significantly affecting all detected PDVs, except for methyl benzoate (Fig. 1e, Supplemental Fig. S2b). This suggests that JA’s mode of action is similar to that of GA, with both downregulating scent production (Ravid et al., 2017). These results are in line with the gradual decrease of JA, and JA-isoleucine and GAs toward anthesis previously observed in petunia flowers (Fonseca et al., 2009; Aravena-Calvo, 2016; Ravid et al., 2017).

### Isolation of EVER- an epidermal MYB-family regulator of volatile emission

The RNA-Seq data generated from the petal adaxial epidermis were used as a platform for the identification of novel genes regulating PDV production. We screened for genes with higher expression in the adaxial epidermis than in the WPT with annotations of TF families in plants (Shiu et al., 2005). This screening yielded 18 genes from 8 distinct TF families: NAC, WRKY, MYB, bHLH, TCP, GRAS, ARF and HD-Zip (Supplemental Table S1). Among them were three MYBs that have been shown to be involved in flavonoid and PDV biosynthesis: *PH4* with a ca. 2.5-fold increase, and *EOBV* and *MYB-FL*, each with a 1.8-fold increase (Quattrocchio et al., 2006; Spitzer-Rimon et al., 2012; Cna’ani et al., 2015; Sheehan et al., 2016); and the GRAS family member Ph*-DELLA* (6.4-fold increase) (Sun, 2011; Ravid et al., 2017). The remaining 14 adaxial epidermis-enriched putative TFs (AETFs) were not previously characterized in petunia. Screening through the blast hits, GO terms and fold-differences of putative MYB AETFs, we chose to concentrate on Peaxi162Scf00777g00521 (*AETF5*). Its expression was 9.1-fold higher in the adaxial epidermis vs. WPT and the best hit of BlastP analysis against Arabidopsis yielded *MYB94* (AT3G47600.1), which has been shown to be enriched in the epidermis and involved in positive regulation of wax biosynthesis (Lee and Suh, 2015). Protein domain analysis of AETF5 using InterPro (Blum et al., 2021) also predicted that it has a MYB-domain at its N terminus. Furthermore, cDNA corresponding to the predicted transcript of *AETF5* was present in our petal MYB cDNA collection generated from corollas of another high PDV-producing *Petunia* x *hybrida* variety, line P720, the same collection that yielded the scent-regulatory MYBs *EOBI*, *EOBII* and *EOBV* (Spitzer-Rimon et al., 2010, 2012); qPCR analysis confirmed the higher expression of *AETF5* in the adaxial epidermis vs. WPT (Supplemental Fig. S1).

To test the relevance of AETF5 to PDV production, a 165-bp fragment of its 5’ UTR was amplified (Supplemental Fig. S3) and cloned into a Tobacco rattle virus RNA2 (TRV2) vector harboring a fragment of *CHS* for transient silencing assays in cv. Mitchell (Fig. 2a) (Spitzer et al., 2007). Localized headspace of volatiles emitted from *AETF5*-silenced petal tissues revealed a significant increase in total emission; increased emission levels were observed for benzyl alcohol, phenylethyl alcohol, vanillin and benzyl benzoate as compared to control pTRV2-*CHS*-inoculated tissues (Fig. 2, b and c). No effect was observed on benzaldehyde, methyl benzoate, phenylacetaldehyde or isoeugenol (Supplemental Fig. S4). Similarly, VIGS of *AETF5* in cv. P720 led to a significant increase in emission of floral PDVs (Supplemental Fig. S5). Taken together, these results suggest that *AETF5* [hereafter *EPIDERMIS VOLATILE EMISSION REGULATOR* (*EVER*)] is involved in negative regulation of PDV emission.

**Figure 2.**
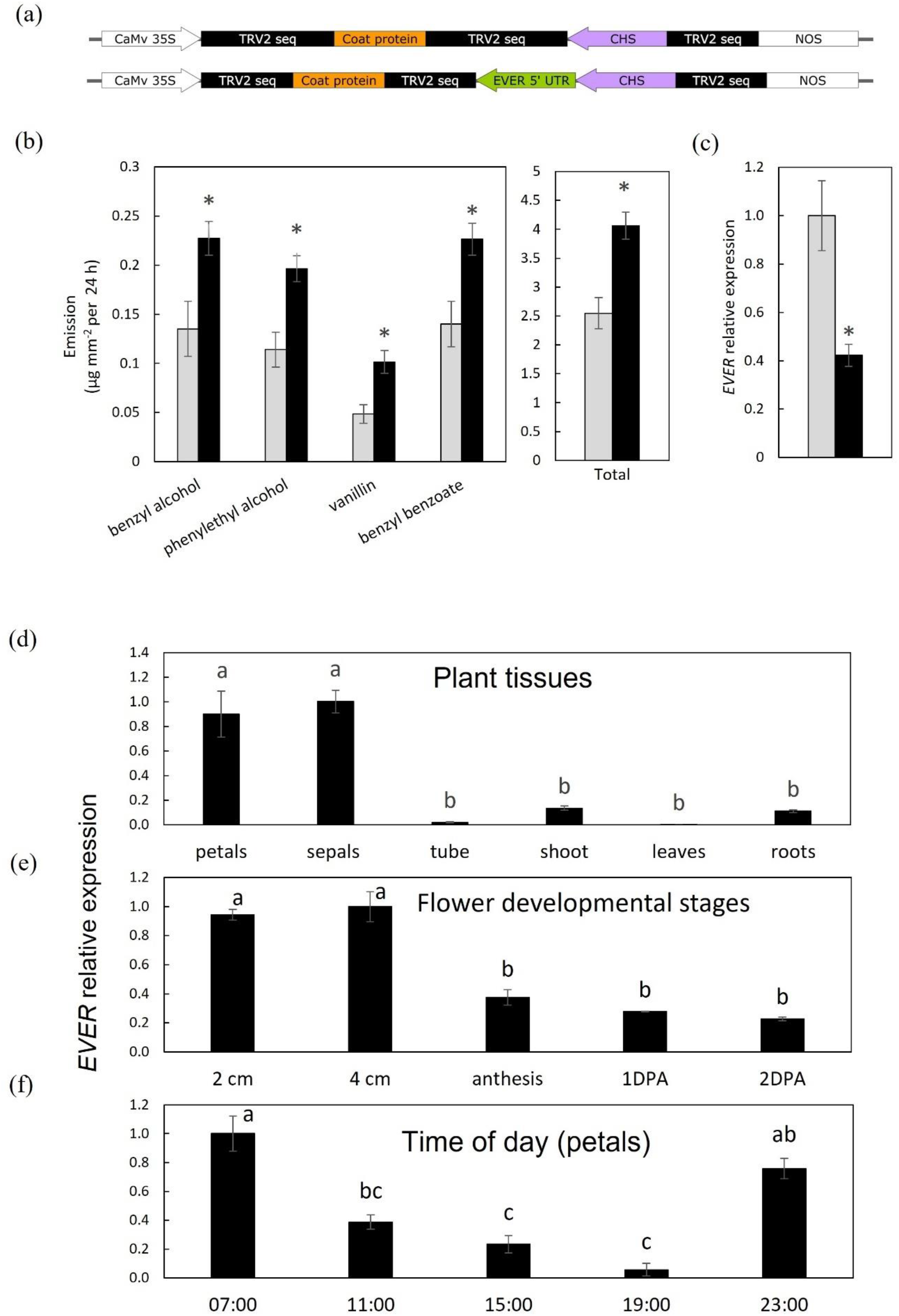
Suppression of *EVER* in petunia ‘Mitchell’ flowers leads to enhanced volatile emission. (a) Schematic representation of the TRV2 vectors used in the experiments. Upper: pTRV2-*CHS*, lower: pTRV2-*CHS-EVER*. (b, c) Flowers were inoculated with agrobacteria carrying pTRV2-*CHS* (gray bars) as a control or pTRV2-*CHS-EVER* (black bars). Flowers were harvested 2 days after inoculation and subjected to localized headspace followed by GC-MS analyses (b) or RNA extraction followed by qPCR analysis (c). Data are means ± SEM (n = 3-7). qPCR data were normalized to *Ubiquitin* and presented data were normalized to those from pTRV2-*CHS*. Significance of differences between treatments was calculated by Student’s t-test: \**P* ≤ 0.05. (d-f) Spatiotemporal expression profile of *EVER.* qPCR data were normalized to *Ubiquitin* and presented data were normalized to sample with highest expression level. Significance of differences was calculated by Tukey’s multiple comparison test following one-way analysis of variance. Values with different letters are significantly different at *P* ≤ 0.05.

### Spatiotemporal and developmental expression profiles of *EVER*

*EVER* expression was analyzed by qPCR and shown to be floral-specific, with highest expression detected in petals and sepals (Fig. 2d). Transcript levels in the flower tube and other examined tissues were significantly lower. *EVER* expression was highest in the petals of young flowers (2 cm and 4 cm buds), declining ca. 2.5-fold at anthesis and then maintaining a steady expression level until 2DPA (Fig. 2e). The diurnal expression pattern of EVER in the petals was rhythmic, with transcript levels peaking in the early morning hours and declining toward midday. Similar expression patterns of *EVER* had been observed in the transcriptome generated from Mitchell buds and flowers, corroborating the qPCR results (Shor et al., 2023b). *EVER*’s developmental expression pattern coincided with the developmentally regulated GA levels (Ravid et al., 2017; Patrick et al., 2021). To test whether GA can induce *EVER* expression, 4 cm buds were treated with GA_3_, and analyzed by qPCR. GA treatment led to a significant increase in *EVER* transcripts compared to control mock-treated flowers (Supplemental Fig. S6). As expected, transcript level of *GIBBERELLIN INDUCED PROTEIN (GIP)* was induced in buds treated with GA, whereas *ODO1* transcript levels dramatically decreased (Ravid et al., 2017).

### *EVER* is an R2R3 MYB activator belonging to family subgroup 1

In the genomes of *Petunia x hybrida* ancestors—the strongly scented *P*. *axillaris* and low-scented *P. inflata*—there is one copy of *EVER* with an atypical intron organization compared to other R2R3-MYBs: it contains four exons and three introns, instead of the common three and two, respectively (Bombarely et al., 2016; Chen et al., 2021). The gene in both ancestors has a predicted uninterrupted open reading frame that produces a functional protein. Protein alignment of EVER showed 95.18% identity with *P*. *axillaris* and 93.89% with *P*. *inflata* (Supplemental Fig. S7).

Although EVER repressed scent emission (Fig. 2b, Supplemental Fig. S4), it does not have an EAR domain like, for example, the negative regulator Ph-MYB4 (Sainz et al., 1997; Colquhoun et al., 2011; Rodrigues et al., 2021). On the contrary, phylogenetic analysis clustered EVER into subgroup 1 of MYB activators (Fig. 3a, Supplemental Fig. S8) (Dubos et al., 2010). Indeed, like other members of this subgroup, it has a conserved R2R3-binding domain at its N terminus that allows interaction with promoter elements, and its C terminus is enriched in acidic amino acids and contains ‘YASS^A^/_T_^E^/_D_NI’ and ‘SL^F^/_I_EKWLF^D^/_E_’ domains which are important for transactivation (Supplemental Fig. S9) (Lee and Suh, 2015). Other notable subgroup 1 members are MYB30, MYB31, MYB94 and MYB96 from Arabidopsis, MYB306 from pepper, and MYBYS from cucumber that were shown to regulate various pathways in plants, such as wax biosynthesis, anthocyanin and carotenoid pathways.(Leitner-Dagan et al., 2006; Lee and Suh, 2015; Lee et al., 2016; Ma et al., 2022; Shi et al., 2022).

**Figure 3.**
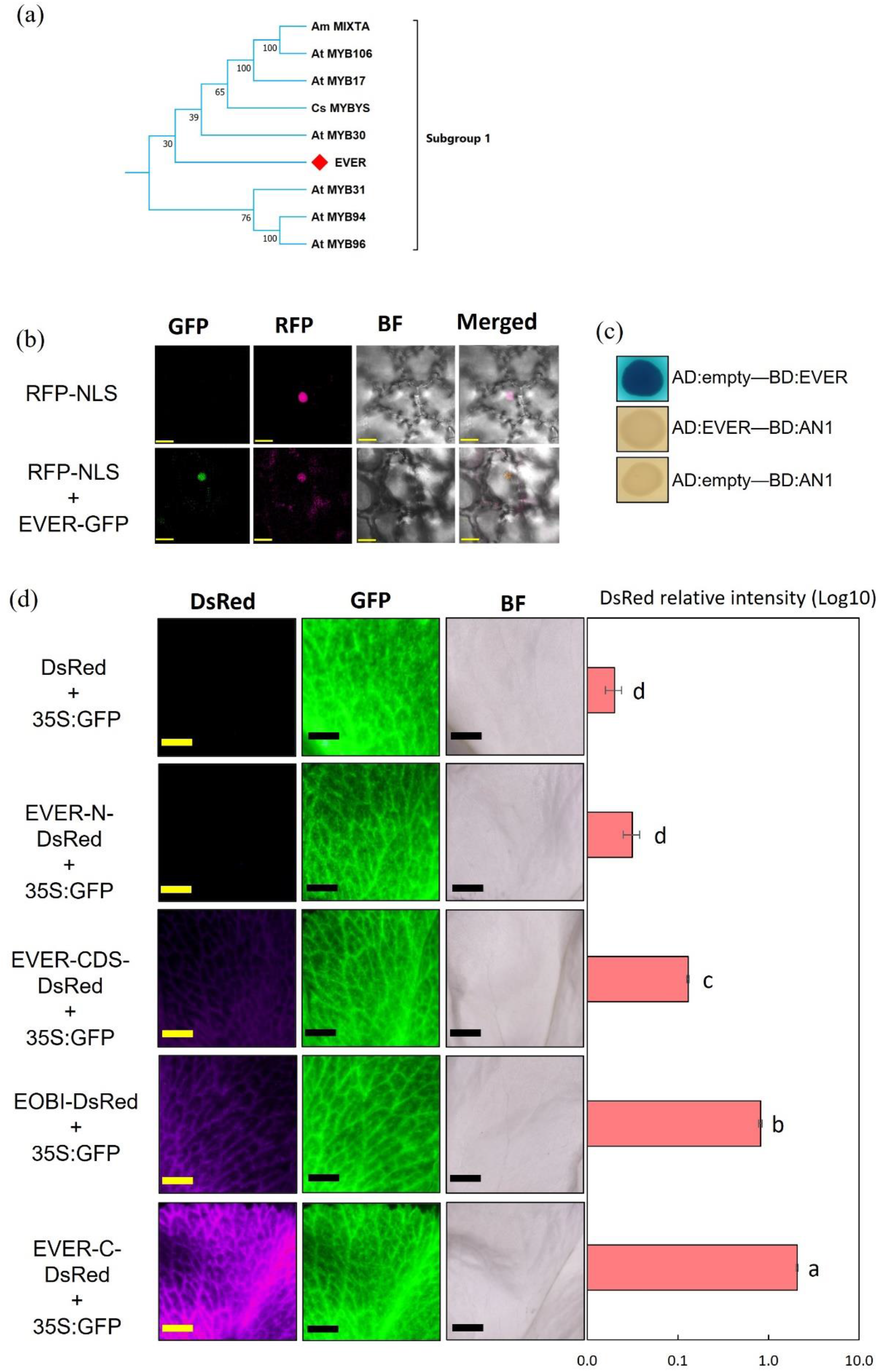
EVER is a nuclear-localized subgroup 1 MYB activator. (a) Branch of phylogenetic tree displaying the similarity between EVER and other R2R3-MYBs belonging to subgroup 1. Protein sequences were aligned by ClustalW, followed by neighbor-joining method with 1000 bootstraps using MEGA version 11. (b) Nuclear localization of EVER in petal epidermal cells revealed by confocal microscopy. Petunia petals were inoculated with agrobacteria carrying EVER-GFP and RFP-NLS or only RFP-NLS as a control. Bars = 10 μm. (c) Transactivation capabilities of EVER in yeast. EVER was fused to GAL4 binding domain (BD) or GAL4 activation domain (control, AD). Yeast transformed with AD:empty and BD:EVER displayed activation of the *LacZ* reporter gene (upper image). In contrast, *LacZ* activation was not observed in yeast transformed with AD:EVER and BD:AN1 (a BHLH protein used as a negative control) or AD:empty and BD:AN1. (d) EVER has transactivation capabilities *in planta*. Petunia petals were inoculated with agrobacteria carrying plasmids with LacI-min35S-DsRed alone or together with either EVER-N (only MYB domain), EVER-CDS (complete coding sequence), EOBI (MYB activator, positive control) or EVER-C (lacking MYB domain). All treatments included agrobacteria carrying 35S:GFP as a reference. Two days after inoculation, flowers were harvested and activation of LacI-min35S-DsRed was examined under a fluorescence stereomicroscope. DsRed and GFP intensities were analyzed using Fiji software. Significance of the differences was calculated by Tukey’s multiple comparison test following one-way analysis of variance. Values with different letters are significantly different at *P* ≤ 0.05. Bars = 3 mm. BF = bright field.

To confirm that EVER is located in the nucleus, it was fused to green fluorescent protein (GFP) and transiently expressed in petunia petals. Confocal microscopy revealed GFP signal in the nucleus together with red fluorescent protein fused to a nuclear-localization signal used as a positive control for nuclear localization (Fig. 3b).

To assess EVER’s ability to transactivate genes, it was fused to the GAL4 binding domain (BD) or activation domain (AD) and tested in a yeast two-hybrid (Y2H) assay. Yeast cells transformed with BD-EVER were intensely stained, indicating strong transactivation. No staining was observed when EVER was fused to AD (Fig. 3c). To test EVER’s transactivation ability *in planta*, and to further characterize its activation domain, its coding sequence (EVER-CDS), C terminus (EVER-C) containing transactivation domains and N terminus (EVER-N) containing DNA-binding domains were fused to LacI-BD (Supplemental Fig. S3). *DsRed* fused to the lac operator–minimal 35S promoter (Lac–min35S), used as a reporter gene for activation, was cloned into each of the above constructs, creating EVER-CDS-DsRed, EVER-C-DsRed and EVER-N-DsRed. The well-characterized floral scent activator EOBI, fused to LacI-BD, was used as a positive control for *in-planta* activation. Petunia petals were inoculated with these constructs together with 35S:GFP (as a reference for inoculation). EVER-C demonstrated the highest activation of DsRed (normalized to GFP)—stronger than EVER-CDS and EOBI (Fig. 3d). In contrast, EVER-N failed to induce DsRed expression, which was similar to the background levels of Lac-min35S-DsRed alone (Fig. 3d). These results demonstrate that EVER is a nuclear-localized MYB with transactivation capabilities.

### Generation of *ever* knockouts by viral-based CRISPR/Cas9 system

To further evaluate EVER’s involvement in the regulation of PDV production, we induced knockout mutations in the gene encoding EVER using the CRISPR/Cas9 system. Gene-editing efficiency is dependent on several factors, including the expression levels of the CRISPR components, which vary among transformation events (Javaid and Choi, 2021). Moreover, in recalcitrant plants, it is extremely difficult to generate multiple transformation, regeneration, and gene-editing events. To overcome this issue, a three-step procedure was devised: (1) *Cas9*-expressing plants were generated; (2) transgenic *Cas9*-expressing plants were assessed for nuclease activity; (3) selected plants were inoculated with pTRV2 carrying sgRNA targeting the desired genes. To this end, *Cas9*-transgenic petunia cv. Mitchell plants were generated by inoculation of explants with *Agrobacterium* carrying *Cas9* under the Arabidopsis ubiquitin 10 promoter, and *DsRED* and *NPTII* genes under the 35S promoter for plant selection (Supplemental Fig. S10a). The presence of *Cas9* in *DsRed*-expressing plants was confirmed by PCR (Supplemental Fig. S10b). To assess the nuclease activity in Cas9-positive lines, they were transiently inoculated with *Agrobacterium* harboring a mutated *uidA* gene (mGUS) and a sgRNA targeting the mutated area within the *mGUS* sequence, followed by GUS histochemical staining (Supplemental Fig. S10, c to e) (Marton et al., 2010). Lines #2 and #55 were selected as the platform for generation of *ever* knockouts using pTRV2 vector carrying sgRNA under a viral subgenomic promoter (Fig. 4a). The genomic sequence of *EVER* in Mitchell was confirmed by Sanger sequencing and the CRISPOR web tool located a suitable sgRNA (sgRNA:EVER) to target the second exon (Concordet and Haeussler, 2018) (Fig. 4b). To allow rapid molecular screening of gene-edited plantlets, the sgRNA was designed to disrupt the recognition sequence of the restriction enzyme *Bsu361* when an indel occurs (Fig. 4b). Almost half of the explants (48%) inoculated with pTRV2-sgRNA:*EVER* generated plantlets, of which 44% harbored mutations in the targeted second exon of *EVER* as revealed by PCR followed by restriction with *Bsu361* (Supplemental Fig. S11). These plants were transferred to the greenhouse, self-pollinated and their progeny molecularly analyzed to confirm inheritance of the mutation and zygosity levels. Following Sanger sequencing, four independent, homozygous lines with frameshift mutations, *ever* #1, 2, 3 and 4, harboring a cytosine deletion, 4-bp deletion, and an adenine or thymine insertion, respectively, were selected for further analysis (Fig. 4b). The four *ever* knockout lines developed and flowered normally, and did not display any observable differences from control, Cas9-expressing plants (Fig. 4b). The described approach has been successfully implemented to induce CRISPR/Cas9-mediated gene editing in other recalcitrant plant species, such as pepper (Supplemental Fig. S12).

**Figure 4.**
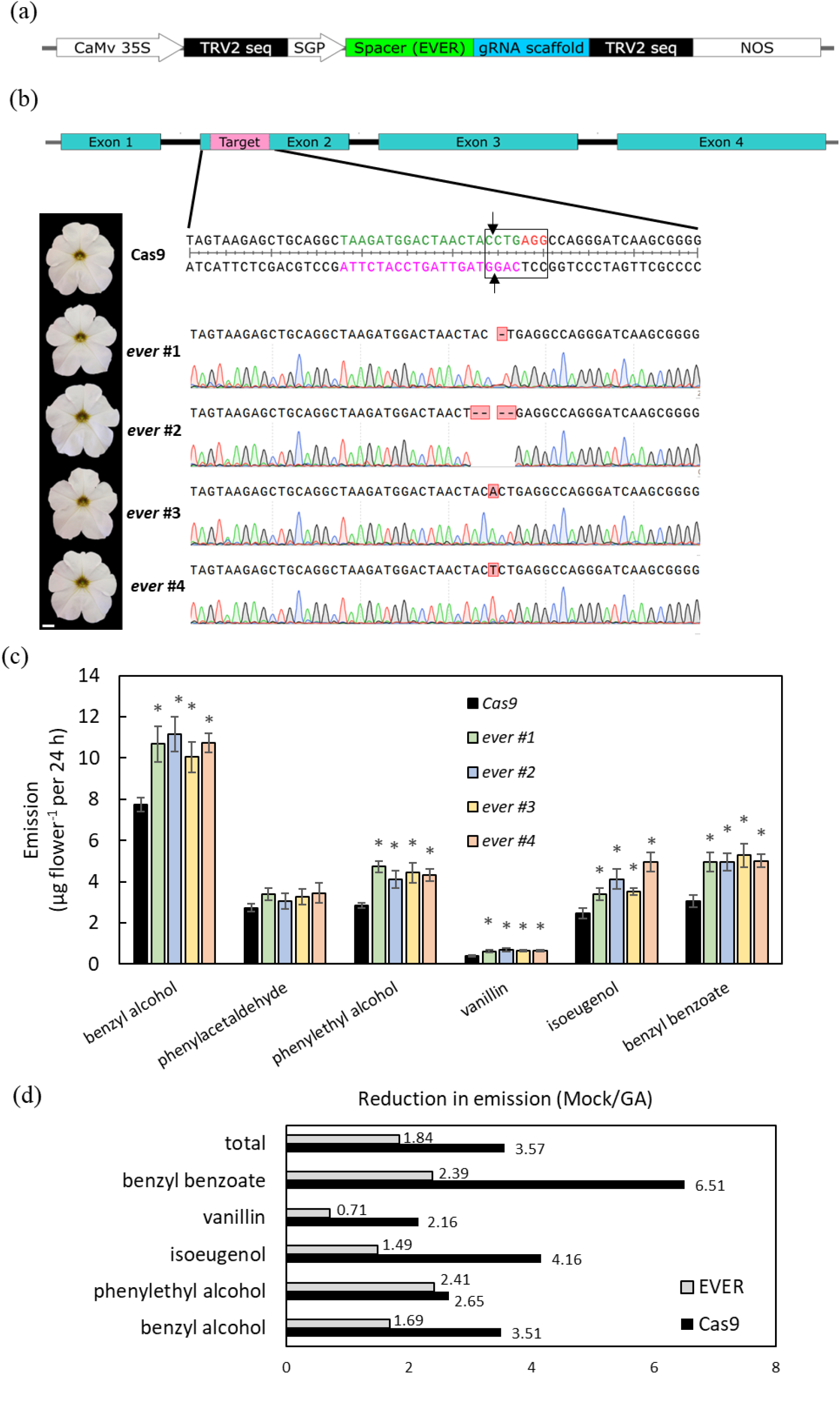
Generation of *ever* knockout lines by viral-based Cas9 system. (a) Schematic representation of the TRV2 vector used for gene editing. (b) Top: Targeted genomic sequence of *EVER*. Green nucleotides, spacer sequence; magenta nucleotides, target; red nucleotides, protospacer adjacent motif (PAM). Black arrows mark the predicted cleavage site of Cas9. Black frame marks the recognition site of *Bsu361* (CCTGAGG) used for molecular screening of mutants. Left: representative 2DPA petunia cv. Mitchell flowers from *Cas9* control and *ever* lines #1-#4. Bar = 1 cm. Bottom right: Sanger sequencing of targeted region in *ever* knockout lines. (c) Dynamic headspace analyses of *ever* knockout lines and *Cas9* control were performed for 24 h on 2DPA flowers followed by GC-MS analyses. Data are means ± SEM (n =15-17). Significance of differences was calculated using Dunnett’s multiple comparison (\**P* ≤ 0.05) with *Cas9* as the control following one-way analysis of variance. (d) Effect of GA on PDV emission is attenuated in *ever* knockouts. Buds of *Cas9* and *ever* #1 lines were treated with or without GA_3_ and after 48 h, flowers were analyzed by dynamic headspace conducted for 24 h, followed by GC-MS analyses. Data are presented as the ratio between mock and GA treatments.

### Increased emission levels of low-vapor-pressure PDVs in *ever* knockouts

To assess the effect of *EVER* knockout on volatiles, dynamic headspace analysis was conducted on the CRISPR-mutated lines. Similar to the aforedescribed results (Fig. 2b, Supplemental Figs. S4 and S5), headspace analyses of the four *ever* lines revealed a significant increase in the emission levels of five low-vapor-pressure PDVs derived from distinct branches of the phenylpropanoid pathway: benzyl alcohol and benzyl benzoate from C6-C1 branch, phenylethyl alcohol (C6-C2), isoeugenol, and vanillin (C6-C3) (Fig. 4c) (Skaliter et al., 2022). Interestingly, the emission of these compounds was recently shown to be highly affected by the cuticle (Liao et al., 2021; Ray et al., 2022). Emission levels of petunia PDVs with higher vapor pressure—benzaldehyde, methyl benzoate and phenylacetaldehyde, as well as the terpenoid α-pinene, were not affected in any of the knockout lines (Fig. 4c, Supplemental Fig. S13a). Internal pool analyses, performed to further detail the effect of *EVER* on PDV production, revealed no significant differences between knockout lines and controls (Supplemental Fig. S13b).

Since EVER’s expression was shown to be induced by GA, a negative regulator of scent (Fig. 2b, Supplemental Fig. S6) (Ravid et al., 2017), we evaluated whether GA’s mode of action is mediated by EVER. Flowers of *ever* knockout and control Cas9 plants were subjected to treatments with GA_3_ followed by dynamic headspace analysis. Both types of GA-treated flowers exhibited a reduction in volatile emissions (Fig. 4d). The effect of GA on low-vapor-pressure compounds in the *ever* knockouts as compared to the control, excluding phenylethyl alcohol, was less drastic (Fig. 4d). GA’s effect on methyl benzoate and benzaldehyde did not differ between *ever* and the control (Supplemental Fig. S13c), suggesting involvement of EVER in GA’s effects on low-vapor-pressure compounds. Ravid et al. (2017) showed that GA repression of volatiles is mediated by DELLA proteins, whose action is based on protein–protein interactions (Livne et al., 2015). To determine whether Ph-DELLA interacts with EVER, Y2H assays were performed. As both EVER and DELLA have autoactivation abilities, a truncated version of the latter—ΔDELLA (residues 188–580)—that does not include the DELLA domain was generated (Gallego-Bartolomé et al., 2012). The Y2H assay revealed no interaction between DELLA and EVER (Supplemental Fig. S13d). These results suggest that while repression of volatiles by GA is partially mediated by EVER, this path probably does not involve an EVER–DELLA interaction.

In an attempt to unravel EVER’s target genes, qPCR analyses were performed on flowers at 2DPA for key scent-related biosynthetic enzymes that were enriched in the transcriptome of the adaxial epidermis. Transcript levels of *CM* and *DAHPS* from the AAA-biosynthesis pathway were unaltered between *ever* lines and the control (Fig. 5a). Similarly, no effect was observed on transcript levels of the rate-limiting enzyme of phenylpropanoid biosynthesis, PAL (Fig. 5a). Furthermore, no significant differences were observed in transcript levels of the dedicated PDV-biosynthesis enzymes whose product emission levels changed in *ever* knockouts: benzoyl-CoA:benzyl alcohol/phenylethanol benzoyltransferase (BPBT), which is responsible for the synthesis of benzylbenzoate, benzyl alcohol and phenylethyl alcohol (Boatright et al., 2004); and phenylacetaldehyde synthase (PAAS), which competes with PAL for synthesis of C6–C2 compounds (Fig. 5a) (Kaminaga et al., 2006; Farhi et al., 2010). Accordingly, there were no differences in expression of the positive regulators *ODO1*, *EOBI* and *EOBII*. Similarly, the expression of Ph*-ABCG12* did not differ between *ever* and control Cas9 plants. As *EVER’s* expression is highest in 4 cm buds (Fig. 2e), qPCR analyses were also performed on RNA extracted from that developmental stage, targeting the transcripts of the same genes as for 2DPA flowers. The results for buds were essentially identical to those obtained for 2DPA (Fig. 5a). Taken together, these results suggest that the effect of EVER on PDV emission is not mediated via transcriptional regulation of the tested scent-related biosynthesis genes. The lack of effect of *ever* knockout on levels of internal pools (Supplemental Fig. S13b) was in accordance with these qPCR results.

**Figure 5.**
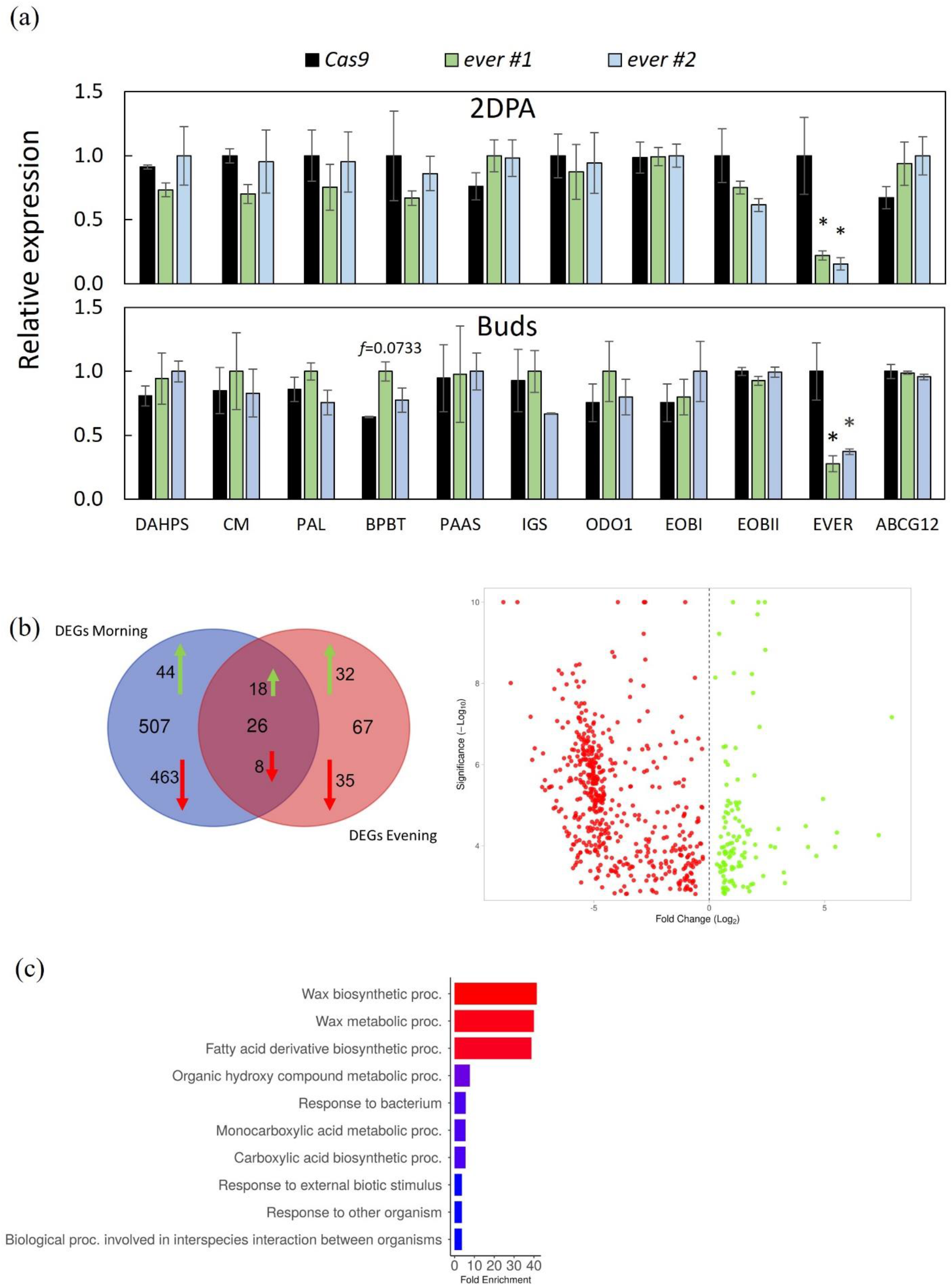
Analyses of gene expression in *ever* knockout lines. (a) qPCR analyses performed on RNA extracted from 2DPA flowers (upper panel) and buds (lower panel) of *ever* lines #1 and #2 and *Cas9* control. Data are means ± SEM (n = 3-5). Data were normalized to the geometric mean of *Ubiquitin* and *EF1α.* Significance of differences was calculated using Dunnett’s multiple comparison (\**P* ≤ 0.05) with *Cas9* as the control following one-way analysis of variance. (b) Venn diagram showing the number of DEGs in *ever* line #*2* in the morning and evening (green arrows represent upregulated genes, red arrows, downregulated genes) (left) and volcano plot of downregulated (red dots) and upregulated (green dots) DEGs in *ever* knockouts vs. controls. (c) GO “Biological process” enrichment analysis of upregulated DEGs in ever knockout line #2.

### Transcriptome of *ever* knockout flowers reveals perturbations in wax-related genes

To gain a deeper understanding of the transcriptomic profile of *ever* knockouts, RNA-Seq analyses were performed on flowers of *ever* line #2 and control Cas9 plants at 1DPA, when scent emission is high, at two time points: morning (1100 h) and evening (1900 h), to identify expression changes that might occur due to rhythmicity (Shor et al., 2023b). The rationale was that *EVER* expression is higher in the morning hours (Fig. 2f), whereas the scent-related machinery is typically active in the evening (Shor et al., 2023b). In the morning samples, 507 exclusive differentially expressed genes (DEGs) (*ever* vs. Cas9) were identified compared to 67 in the evening; 26 genes overlapped between the two samples (Fig. 5b, Supplemental Data S3 to S5). Most DEGs were downregulated, with only ca. 20% upregulated. Genes tested earlier in the qPCR analysis: *CM*, *DAHPS*, *PAL*, *PAAS IGS* and *BPBT*, were not differentially expressed, in line with the qPCR results (Fig. 5a). GO enrichment analysis of “Biological process” performed on all DEGs, i.e., downregulated and upregulated in morning and evening, yielded general pathways (Fig. S14a). Since most DEGs in both morning and evening datasets were downregulated (Fig. 5b), the GO analyses were biased toward them, as seen by enrichment conducted only on downregulated DEGs (Fig. S14b). Therefore, GO analyses were performed on upregulated morning and evening DEGs only. The analysis showed enrichment in “Wax biosynthetic process”, “Wax metabolic process”, and “Fatty acid derivative biosynthetic process” for “Biological process” (Fig. 5c). Among these enriched genes were *CYCLOPROPANE-FATTY-ACYL-PHOSPHOLIPID SYNTHASE* (Peaxi162Scf00266g01010), *LACS* (Peaxi162Scf00786g00434), *FATTY ACID HYDROXYLASE SUPERFAMILY* (*CER3*), *AP2/B3-like TRANSCRIPTIONAL FACTOR FAMILY PROTEIN* (Peaxi162Scf00420g00827) and *FATTY ACYL-COA REDUCTASE 2* (Peaxi162Scf00463g00320) (Supplemental Data S5 andS6).

To gain further information on EVER’s mode of action, transient overexpression of *EVER* was performed in petals followed by RNA-Seq analysis. Ectopic expression of *EVER* resulted in enrichment of genes involved in “Wax biosynthetic/metabolic process” and “Fatty acid derivative biosynthetic process” (GO Biological process), in line with the results obtained for the transcriptome of *ever* knockouts (Fig. 5c, Supplemental Fig. S15). As expected, *EVER* was significantly overexpressed in petal tissues inoculated with 35S:EVER as compared to the DsRed control (Supplemental Data S6). Transcripts of genes that are targets of MYB94 and MYB96 in Arabidopsis (homologs of EVER) were also upregulated in 35S-EVER petunia corollas (Lee and Suh, 2015; Lee et al., 2016): Peaxi162Scf00087g00513 (*3-KETOACYL-COA SYNTHASE 1*), Peaxi162Scf00104g00816 (*3-KETOACYL-COA SYNTHASE 6*), Peaxi162Scf00420g00236 (*ESTRADIOL 17-BETA-DEHYDROGENASE 12*) and Peaxi162Scf00314g00943 (*CER3*) were also upregulated. In addition, Peaxi162Scf00034g00413 (Ph-*AA13*), shown to encode an enzyme that converts malonic acid to the precursor of epicuticular waxes malonyl-CoA, was also upregulated in EVER-overexpressing tissues (Chen et al., 2017).

### The petals of *ever* knockouts display altered composition of epicuticular petal waxes

We further explored the possible involvement of EVER in wax biosynthesis in petunia flowers, having obtained the following pieces of evidence suggesting this link: (1) the homologs of EVER in Arabidopsis were shown to regulate the biosynthetic pathways of waxes; (2) our RNA-Seq data of *EVER*-knockout and overexpressing lines revealed enrichment of genes involved in wax-biosynthesis processes. We extracted the epicuticular waxes from 2DPA petals of *ever* #1 and #2 and control Cas9 lines and profiled their composition via GC–MS-based protocols (Sarkar et al., 2023). We positively identified 29 wax-related components, most of which belonged to wax esters previously shown to dominate epicuticular waxes in petunia flowers (King et al., 2007; Chen et al., 2017). The most abundant esters were C20 hexyl, phenylmethyl and 3-methylbutyl (Supplemental Data S7). Comparisons of the epicuticular wax compositions revealed significantly lower levels of several esters and fatty acids in petals of *ever* #1 and #2 compared to Cas9 control lines, including the esters C18 phenylmethyl, C22 hexyl, C22 *n*-octyl, C24 hexyl and isovaleric acid eicosyl, and fatty acids C18 and C20 (Fig. 6). Reductions in the latter two fatty acids were also detected in analyses of the wax by LC–MS (Supplemental Fig. S16 and Data S7). These reductions resulted in significant decreases in total wax loads in petals of both mutant lines (Fig. 6). The ester C18 phenylmethyl was the only component that was at significantly higher levels in flowers of both *ever* knockout lines compared to the Cas9 control (Fig. 6).

**Figure 6.**
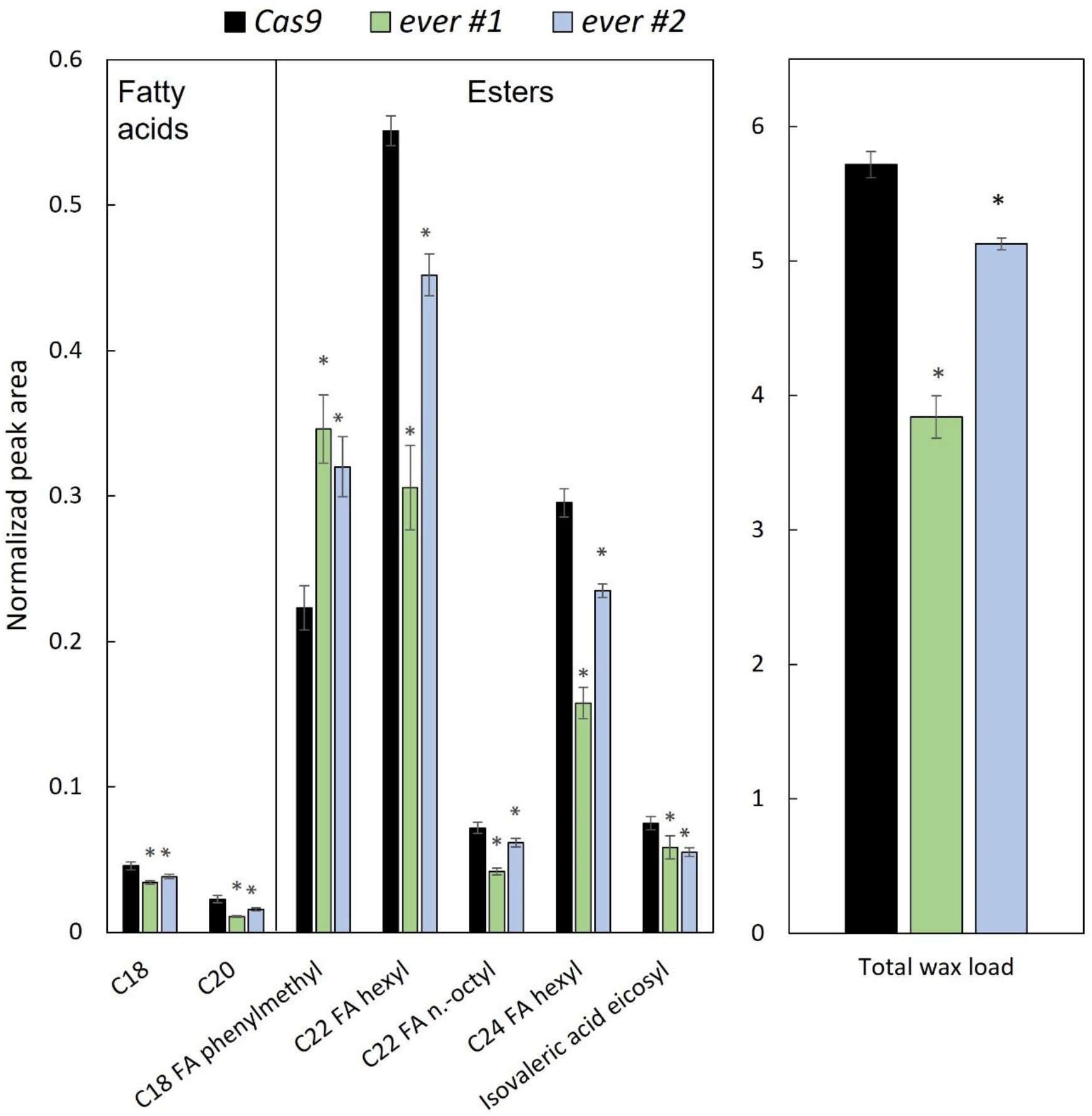
Epicuticular wax composition and load are altered in petals of *ever* knockout lines. Wax was extracted from petals of *ever* lines #1 and #2 and the *Cas9* control followed by GC-MS analysis. Data are means ± SEM (n = 4). Significance of differences was calculated by Dunnett’s multiple comparison (\**P* ≤ 0.05) with Cas9 as the control following one-way analysis of variance.

Downregulating Ph*-ABCG* transporter(s) in petunia petals has been shown to compromise the permeability of the cuticle, leading to abnormal epidermal cells and flowers (Liao et al., 2021). In this study, we did not detect any differences in the phenotype of *ever* knockout flowers or epidermal cells, as revealed by cryogenic scanning electron microscopy (SEM) (Fig.4b and Supplemental Fig. S17, a to i). Furthermore, flowers of the knockout lines looked similar to controls following toluidine blue staining, and their rate of water loss and cuticle thickness was practically the same (Supplemental Fig. S17, f to k and Fig. S18 a and b). These results indicate that the alterations in wax composition in petals of *ever* knockout lines did not compromise the cuticle. As *EVER* expression was specific to flowers, we speculated that the reductions in epicuticular waxes detected in *ever* flowers would not occur in leaves. To validate this, we profiled the epicuticular wax composition of ever #1 and #2 and control Cas9 leaves via similar GC–MS protocols described above. The composition was far less rich than that the one detected in petals and was dominated by alkanes, particularly of C31 alkane (Supplemental Data S8). As expected, the levels of all 11 identified epicuticular waxes in leaves were identical in *ever* knockout and control leaves (Supplemental Fig. S18 and Data S8). Taken together, our results provide solid evidence that EVER is involved in the biosynthesis of epicuticular waxes in petunia petals.

## Discussion

To establish a platform for the identification of spatially enriched genes in petunia flowers, we generated RNA-Seq data from the epidermis of the petal of petunia cv. Mitchell. We chose to examine the adaxial epidermis as it has been shown to emit significantly higher levels of volatiles, and to have higher expression of scent-related genes, than the abaxial layer (Skaliter et al., 2021). Transcriptomics has proven to be a powerful tool in scent studies in petunia for the identification of novel genes (Liao et al., 2021, 2023; Shor et al., 2023b), but the transcriptomes in those studies were generated from whole petal tissues that include all petal layers. Although it is technically simpler to extract RNA from homogenized tissues, and indeed novel genes have been identified using those transcriptomes, spatial differences in transcript expression may result in loss of data, such as dilution of epidermis-enriched transcripts with low expression, leading to missing potential genes. For example, it was reported that 58 genes that were highly expressed in epidermal layers of *Medicago truncatula* leaves, whereas their expression was extremely low in the whole leaf (Cui et al., 2022). Similarly, in the current research, 178 genes displayed expression levels that did not pass the minimum cutoff (>30 max count reads) in the WPT, but did pass the cutoff in the adaxial epidermis—*EVER* being among them.

Interestingly, RNA-Seq revealed that most of the known scent-related genes (except for *CM1*, *DAHPS1, EOBV*, *PAL* (a and c) and *CFAT*) were not preferentially expressed in the adaxial epidermis. However, this cannot be explained by tissue contamination, because microscopic observation of tissue extracts (Skaliter et al., 2021) and expression levels of genes with previously characterized expression patterns [*PH3* and *PH5*, *LACS*, *PDF2*, *EP1*, *FBase*, *CAB* and *PH4* (Supplemental Data S2, Fig. S1)] authenticated the experimental procedure. This leads us to suggest that most of the genes involved in phenylpropanoid scent-related pathways are not expressed exclusively in the epidermis. Several lines of evidence support these observations. For example, it has been demonstrated that promoters of scent-related genes are also active in mesophyll cells of petunia petals (Benfey et al., 1990; Van Moerkercke et al., 2012). Interestingly the major rate-limiting enzymes DAHPS, CM and PAL are epidermis-specific, suggesting that the metabolic flow into the phenylpropanoid pathway is still epidermally controlled. In this respect, it would be informative to analyze metabolic fluxes within each layer of the petal.

To detail the functionality of EVER, we generated *ever* knockout lines using a three-step viral-based CRISPR/Cas9 system. Use of the CRISPR/Cas9 system has been reported in *P*. *axillaris* (Lüthi et al., 2022) and *P. hybrida* (Chopy et al., 2020; Lin and Jones, 2022). Here we implemented a viral vector to deliver sgRNA to petunia tissues, expanding the existing toolkit for efficient gene editing. The viral-based system used in this research presents some advantages compared to “traditional” Cas9 systems. The preliminary “optimal nuclease activity” screening allows eliminating plants with very low Cas9 activity, thus saving and improving efficiencies. The viral delivery of sgRNA has four advantages: (1) the virus’ ability to spread throughout cells of the inoculated tissue increases the chance for mutations in cells that were not infected with agrobacteria as well; (2) as the sgRNA is driven by a viral promoter, and not the commonly used *U6* promoter, there is no prerequisite for a guanine nucleotide start site in the spacer sequence, thus allowing more flexibility in target selection (Long et al., 2018); (3) several sgRNA can be cloned under the same subgenomic promoter allowing shorter T-DNA and requiring fewer cloning steps; (4) the virus-driven sgRNA activity decreases with plant development and this reduces the chances of off-target mutations. In addition, plantlets are regenerated without the use of antibiotic-selective conditions which again, improves yield, especially where recalcitrant plants, such as pepper, are concerned (Supplemental Fig. S12).

Knockout lines, localized transient assay and VIGS revealed an increase in volatile emission in *EVER*-suppressed tissues (Figs. 2b and 4c, Supplemental Fig. S5). The effect on emission was not exclusive to one main branch of PDV biosynthesis: benzyl alcohol and benzyl benzoate (C6–C1 branch), phenylethyl alcohol (C6–C2) and isoeugenol and vanillin (C6–C3) all demonstrated enhanced emission. In contrast to these low-vapor-pressure compounds, the emission of volatiles with higher vapor pressure, e.g., benzaldehyde, phenylacetaldehyde and methyl benzoate, was not significantly affected in flowers with suppressed expression of *EVER*. While in many instances, emission levels and internal pools of volatiles and expression of scent-related genes are in direct correlation (for review see Skaliter et al., 2022), in *ever* knockout lines, internal pools and transcript levels of known scent-related genes—in contrast to emission levels—were unaltered. This phenomenon has been documented previously: transgenic roses ectopically expressing PRODUCTION OF ANTHOCYANIN PIGMENT1 from Arabidopsis displayed increased emission levels of the norisoprenoid β-ionone (Zvi et al., 2012). Nevertheless, similar to *ever* lines, internal pools were unchanged and transcript level of its corresponding enzyme CLEAVAGE DIOXYGENASE1 was downregulated. The machinery coordinating volatile production, i.e., biosynthesis, storage and transport within and outside the tissue still needs to be explored.

The developmental expression pattern of *EVER* was negatively correlated with the production of PDVs—decreasing significantly toward onset of scent production. This pattern is atypical compared to other identified MYB scent regulators, including negative regulator Ph-MYB4 (Colquhoun et al., 2011). EVER’s mRNA profile coincided with that of GA accumulation, and a connection between GA and wax has been demonstrated in tomatoes; a mutation in *FIRM SKIN 1*, encoding a GA2-oxidase, resulted in increased bioactive GA content with concomitant enhanced wax biosynthesis (Li et al., 2020). The link between GA and EVER was demonstrated here by external application of GA_3_, which significantly enhanced *EVER* expression, and by the fact the GA’s negative effect on volatiles was attenuated in the *ever* lines (Fig. 4d). Whereas no protein interaction between DELLA and EVER was observed in the Y2H assay (Supplemental Fig. S13d), we cannot discount the fact that GA’s effect on EVER may be exerted via DELLA’s transactivation abilities. Alternatively, DELLA-independent pathways may be responsible for GA’s activation of *EVER* (Livne et al., 2015). Similar to the negative effect of GA on scent emission, based on gene ontology, we revealed JA’s involvement in floral scent emission. JAs and GAs seem to behave similarly; their concentrations are high in buds and decline toward flower opening, which is in line with their similar effects on scent production. Evaluating the interplay between these two hormones in petunia corollas will shed light on developmental control of scent production.

Examination of scent-related transcripts in CRISPR-induced *ever* knockout plants did not reveal genes that are affected by EVER. On the other hand, GO analyses indicated changes in the expression of genes involved in wax formation. LC/GC–MS analyses of epicuticular wax extracted from petals revealed alterations in its fatty acid and ester levels and ratios. These changes in petal wax composition may explain the increased emission of low-vapor-pressure PDVs observed in the *ever* lines. Liao et al. (2021) showed that compromised cuticle, i.e., a significant decrease in the levels of all wax components and in its thickness, with abnormal flowers and flat epidermal cells, affects low-vapor-pressure volatiles’ emission rates due to the mass-transfer limitations of the cuticle. Wax in *ever* knockouts was reduced by only ca. 20% and this did not affect plant/flower development, shape of the epidermal cells or cuticle permeability. While total wax load was reduced, levels of some of wax components increased, leading to a significant modification in epicuticular wax composition. A study using a model cuticle mimicking petunia floral wax was conducted to elucidate the effect of wax-composition changes on diffusion/emission of volatiles (Ray et al., 2022). Those authors suggested that the ratios between different cuticular wax components affect its structure and therefore, the diffusion rate of different volatiles, especially those with high molecular weight. For example, their results showed that when wax was enriched in 3-methylbutyl dodecanoate, a fatty acid ester, diffusivity of phenylethyl alcohol and benzyl benzoate was highest, whereas it was lowest when a fatty alcohol (1-docosanol) was the main component (Ray et al., 2022). On the other hand, modifying resistance barriers has had varying effects on emission, internal pool levels and expression of scent-related genes. For example, downregulation of a non-specific lipid transfer protein which facilitates transportation of volatiles through the cell wall inhibited PDV emission in petunia flowers, without affecting internal pools or scent-related genes, but rather the volatiles’ levels in the wax. Suppression of *PH4* or *ABCG1* (a plasma membrane transporter) resulted in a reduction in PDV emission levels with a concomitant increase in internal pools; no effect on transcripts of scent-related genes was reported (Cna’ani et al., 2015; Adebesin et al., 2017). Overexpression of *ABCG1* in the abaxial, but not adaxial epidermis led to enhanced emission levels, whereas *PH4* overexpression did not have any effect (Skaliter et al., 2021). A detailed characterization of petunia genes responsible for protective barrier formation, most of which are not annotated, should enable the elucidation of its precise effect on the mechanism of floral scent production. The identification of the transcriptional regulator EVER, which affects the emission of volatiles via modifications of epicuticular wax, should be a valuable tool in deciphering the steps interconnecting the scent-production machinery.

## Materials and methods

### Plant material and growth conditions

*Petunia × hybrida* cv. Mitchell diploid seeds were kindly provided by Cai-Zhong Jiang, University of California, Davis, USA. Rooted plants of *Petunia × hybrida* cv. P720 were obtained from Danziger – “Dan” Flower Farm (Mishmar Hashiva, Israel). Plants were grown in a glasshouse under 25 °C day/20 °C night temperatures with a 16 h light/8 h dark photoperiod.

### RNA extraction and qPCR analyses

Total RNA was extracted from 30–100 mg of ground (with liquid nitrogen) petal tissues using the Tri-Reagent Kit (Sigma-Aldrich, https://www.sigmaaldrich.com) and treated with RNase-free DNase I (ThermoFisher Scientific, https://www.thermofisher.com). First-strand cDNA was synthesized using 1 μg of total RNA, oligo(dT) primer and reverse transcriptase ImProm-II (Promega). Two-step quantitative real-time PCR(qPCR) was performed on a CFX Opus 384 Real-Time PCR System (Bio-Rad, https://www.bio-rad.com/) using 2X qPCRBIO SyGreen Blue Mix Hi-ROX (PCR Biosystems, https://pcrbio.com). A standard curve was generated for each gene using dilutions of cDNA samples, and data analysis was performed using rotor-gene q 2.1.0. Primer specificity was determined by melting-curve analysis. The primers are shown in Supplementary Table S2.

### Generation of the transcriptomic databases

Samples from ‘Mitchell’ flowers were collected 1DPA at 1900 h. Epidermis was isolated as described in Skaliter et al. (2021). RNA extraction was performed using RNeasy Plant Mini Kit (Qiagen, https://www.qiagen.com) according to the manufacturer’s instructions, followed by DNase treatment using Invitrogen’s DNA-free kit DNase. First-strand cDNA was synthesized using total RNA, oligo(dT) primer and reverse transcriptase ImProm-II (Promega, https://www.promega.com). Three biological replicates for each type of sample were used for sequencing. 100 bp single-end reads were sequenced on Novaseq SP of an Illumina NovaSeq. The output was ∼21 million reads per sample. Reads were mapped to the Petunia axillaris reference genome (https://solgenomics.net/ftp/genomes/Petunia_axillaris/Peaxi162annotation_v4.gff). Libraries were sequenced as in Shor et al. (2023b).

### Suppression of *EVER* in petunia flowers via VIGS

*Agrobacterium tumefaciens* strain AGLO was transformed with plasmids harboring RNA1 (pTRV1), pTRV2-*CHS-EVER* or pTRV2-*CHS*. Agrobacteria carrying pTRV1 were mixed in a 1:1 ratio (to OD_600nm_ 0.5 or 10 for transient suppression or VIGS, respectively) with agrobacteria carrying either pTRV2-*CHS-EVER* or pTRV2-*CHS* in inoculation solution containing 200 µM acetosyringone and 10 mM MgCl_2_. For transient suppression of *EVER*, corollas of ‘Mitchell’ flowers at anthesis were inoculated with the *Agrobacterium* solutions by piercing with a needle and infiltrated using a syringe, followed by localized headspace analysis as in Skaliter et al. (2021). For VIGS assay in line P720, *Agrobacterium* solutions were applied to cut stems (after removing apical meristems). After a month, flowers with white areas (indicating viral spread) were used for volatile analyses.

### Collection of emitted volatiles, internal pools and GC–MS analysis

Dynamic headspace analysis was performed by harvesting flowers 2DPA and placing them in a 50 mL beaker filled with tap water in jars (2 flowers per jar). Localized headspace analysis was conducted as in Skaliter et al. (2021). Volatiles were collected for 24 h using columns made of glass tubes containing 100 mg Porapak Q polymer and 100 mg 20/40-mesh, held in place with plugs of silanized glass wool. Trapped volatiles were eluted with 1.5 mL hexane and 0.5 mL acetone. Isobutyl benzene was used as an internal standard.

To determine the pool sizes of volatile compounds, 200 mg tissue from 2DPA flowers was ground in liquid nitrogen and extracted in 400 μL hexane containing isobutylbenzene as the internal standard. Following 2 h of incubation with gentle shaking at 25 °C, extracts were centrifuged at 10,500*g* for 10 min. GC–MS analyses of volatile organic compounds were performed as described in Skaliter et al. (2021).

### EVER localization assay

The CDS of *EVER* (without the stop codon) was amplified from cDNA extracted from flowers and fused to GFP using a GoldenBraid reaction (Sarrion-Perdigones et al., 2013) to generate pDGB3α2-EVER-GFP. ‘Mitchell’ flowers at anthesis were inoculated with a mix of agrobacteria carrying pDGB3α2-EVER-GFP and pRCS-RFP-VirD2-NLS (marker for nuclear localization) or pRCS-RFP-VirD2-NLS alone as a control. Inoculated flowers were harvested 2DPA and imaged in a Leica TCS SP8 confocal system (Leica Microsystems, https://www.leica-microsystems.com) with LAS X Life Science Microscope software. Merged images were generated using Fiji (Image J) software (Rueden et al., 2017).

### EVER transactivation assays

Y2H: This experiment was performed as previously described (Wiseglass et al., 2019). Briefly, the CDSs of the tested genes were cloned to either the GAL4 DNA-binding domain in pBD-GAL4 vector or GAL4-activating domain in pACT2 and transformed into *Saccharomyces cerevisiae* strain Y190. Transformed yeast were grown on synthetic defined medium lacking leucine and tryptophan for selection, followed by X-Gal staining for detection of *β*-*galactosidase* activity.

*In planta*: All vectors were constructed using GoldenBraid (Sarrion-Perdigones et al., 2013). DsRed CDS was cloned under the 6xLacI Operon fused to CaMV 35S minimal promoter (pDGB3α1-2pOpLacI:mini35s-DsRed). GFP cloned into pDGB3α2 under a 35S promoter was used for normalization. EOBI CDS, EVER CDS, and EVER N terminus (EVER-N, amino acids 1–116) and C terminus (EVER-C, amino acids 117– 315) were fused with the LacI binding domain into pDGB3α2. Two days after inoculation, flowers were harvested and examined under a fluorescence stereomicroscope (Nikon SMZ1270, https://www.nikon.com/). DsRed and GFP intensities were analyzed using Fiji (Image J) software (Rueden et al., 2017).

### Generation of *Cas9*-transgenic plants and knockouts

Human-optimized Cas9 (hCas9) was cloned under the control of the *Arabidopsis thaliana* ubiquitin promoter together with *NPTII* under the NOS promoter and DsRed under the 35S promoter using the GoldenBraid 4.0 reaction (Sarrion-Perdigones et al., 2013) to generate pDGB3α1-AtUbqp(1460):hCas9/Nosp:NPTII, which was transformed into *Agrobacterium tumefaciens* strain AGLO. Sterile petunia ‘Mitchell’ explants (young leaves) were incubated for 20 min in a suspension of *Agrobacterium* (in Luria-Bertani broth) conatining 100 µM acetosyringone and the recombinant plasmid pDGB3 pDGB3α1-AtUbqp:hCas9/Nosp:NPTII/35S:DsRed. Explants were placed in the dark for 3 days on Murashige and Skoog (MS) medium supplemented with 3% (w/v) sucrose, followed by transfer for selection on MS with 3% sucrose, antibiotics (kanamycin, carbenicillin), hormones (1.5 mg L^−1^ BA, 0.15 mg L^−1^ NAA), and left under a light/dark 12 h/12 h photoperiod at 22 °C until regeneration and elongation of regenerants were observed. The medium was renewed every 2 weeks. At the elongation stage, BA concentration was decreased to 1 mg L^−1^. To switch regenerants to rooting, they were transferred to media without BA, supplemented with 0.25 mg L^−1^ NAA. The regenerated plants (T0) were tested for DsRed fluorescence, indicating the presence of active introduced genes. Presence of *Cas9* was confirmed by PCR on gDNA using primers 5’-TGGAGGAGTCCTTTTTGGTG-3’ and 5’-GCTTTGGTGATCTCCGTGTT-3’. Positive plants were grown on MS with 3% sucrose, then transferred to the soil. For Cas9 activity test, pAGM-U6p:gmGUS harboring the spacer targeting the premature stop codon of the engineered uidA gene (mGUS) was constructed using Golden Gate and construction of pRCS-p2X35S:mGUS was as previously described (Honig et al., 2015). Leaves of confirmed *Cas9*-transgenic plants were co-infiltrated with *Agrobacterium* harboring pAGM-U6p: gmGUS and pRCS-p2X35S: mGUS or pRCS-p2X35S:mGUS as a control, followed by GUS histochemical assay. Petunia plants demonstrating high Cas9 activity were self-pollinated and T1 plants were tested again for GUS activity and self-pollinated to obtain T2, followed by progeny testing to identify Cas9-homozygous lines. T3 homozygous lines were used as the platform for introduction of *Agrobacterium* harboring pTRV2-sgRNA:*EVER* (synthesized by Twist Bioscience http://www.twistbioscience.com/) and pTRV1. Generated calli were tested for the presence of mutations in *EVER* by PCR (the primers are shown in Supplementary Table S2), followed by restriction analysis with *Bsu361* (ThermoFisher Scentific). Positive results were confirmed by Sanger sequencing, and these plants were further grown and self-pollinated to obtain homozygous mutated lines. T2 generation knockouts were used for experiments.

### GUS and toluidine blue O (TBO) staining assays

GUS staining was performed as previously described (Skaliter et al., 2019). Toluidine blue staining was performed as in Liao et al. (2021). Flowers were collected 2DPA at 1200 h and submerged in 0.05% (w/v) toluidine blue aqueous solution for 3 h, then carefully rinsed with distilled water to remove excess solution.

### Epicuticular wax extraction and analysis

GC–MS: Epicuticular waxes were extracted from 11 mm discs of 2DPA flowers or leaves of *ever* knockout lines and Cas9 control lines by immersion in GC–MS-grade chloroform containing C36 alkane as internal standard (1mg µL^−1^ chloroform). Chloroform extracts were then treated, derivatized and analyzed by GC–MS as recently described by Sarkar et al. (2023).

LC–MS: Flowers of control Cas9 and *ever* lines #1 and #2 were collected at 2000 h and excised using a scalpel to separate them from the tube, gynoecium and androecium. Waxes were extracted from excised corollas of five flowers agitated in 20 mL HPLC-grade hexane for 30 s. The solvent was decanted into a fresh glass vial and left in a chemical hood until completely dried. LC–MS was performed as previously described (Wu et al., 2018). Briefly, wax residue was resuspended in 70% (v/v) acetonitrile and 30% (v/v) isopropanol. LC–MS data were obtained using a Waters Acquity UPLC system equipped with a C8 reverse-phase column (Waters, https://www.waters.com), coupled to a q-Exactive mass spectrometer (Thermo Fisher). Capillary voltage was set to 3 kV, with a sheath gas flow value of 60 and an auxiliary gas flow of 20 (arbitrary units). Chromatograms from the UPLC /MS runs were analyzed and processed with REFINER MS 11.0 (GeneData).

### Scanning electron microscopy

For cryo-SEM and TEM analyses, flowers were collected 2DPA and 3 mm discs were excised. Cryo-SEM samples were mounted with OCT cryo-sectioning media on a flat specimen holder and cryo-fixed by immersion in a nitrogen slush using a Gatan Alto 2500 Cryo-Preparation System (https://www.gatan.com). The microphotographs are recorded using scanning electron microscope JEOL (https://www.jeol.com/) model, JSM-7800F at −140C. TEM samples were fixated in glutaraldehyde 2% and formaldehyde 2% in n 0.1 M sodium cacodylate buffer at room temperature overnight. Samples were rinsed again in cacodylate buffer 3 times (every 10 min) and postfixed in 2% osmium tetroxide overnight. Samples were dehydrated with a graded series of ethanol and embedded in EMbed-812 resin. The microphotographs are recorded using scanning transmission electron microscope detector (Deben, UK) on a JEOL (https://www.jeol.com/) HRSEM, JSM-7800F.

### Phylogenetic analysis

Protein sequences were aligned using the Clustal W algorithm (Larkin et al., 2007) implemented in MEGA11 (Tamura et al., 2021). Phylogenetic analysis was conducted in MEGA11 with default parameters and 1000 bootstraps.

### Statistical and bioinformatic analyses

Statistical analyses were performed using JMP Pro 16 (SAS). Volcano plots were generated using VolcaNoseR (https://huygens.science.uva.nl/VolcaNoseR/). GO and KEGG enrichment analyses were performed using ShinyGO 0.77 (http://bioinformatics.sdstate.edu/go/).

## Data availability statement

Raw RNA-Seq reads were deposited at the National Center for Biotechnology Information (NCBI) under accession number PRJNA971370.

## Acknowledgements

We thank Danziger – “Dan” Flower Farm for providing the plant material. We also thank Dr. Einat Zelinger and Dr. Tally Kossovsky from the Interdepartmental Equipment Unit (Hebrew University of Jerusalem, Rehovot, Israel) for help with the cryogenic electron microscopy. This work was supported by the Israel Science Foundation (grant no. 2511/16). Work in AV’s laboratory is supported by the Chief Scientist of the Israel Ministry of Agriculture and Rural Development (no. 20-01-0209) as part of the National Center for Genome Editing in Agriculture. AV is an incumbent of the Wolfson Chair in Floriculture.

## Author contributions

OS planned and designed the research, performed the experiments, analyzed the data, and wrote the manuscript. DB, E. Shor, E. Shklarman, EM, JA-C, SK, AC, TM, GD, OE and BR performed the experiments and analyzed the data. WJ and YB performed the LC–MS experiment and analyzed the data. HC analyzed the data and wrote the manuscript. AV planned and designed the research and wrote the manuscript. All authors revised the manuscript and approved the final version.

## Supplemental data

**Supplemental Figure S1. Quantitative real-time PCR analyses of RNA extracted from adaxial epidermal tissue and whole petal tissue of *Petunia x hybrida*.**

**Supplemental Figure S2. Enriched GO “Biological process” terms in whole petal tissue.**

**Supplemental Figure S3. mRNA sequence of *EVER* (*AETF5*) from *Petunia x hybrida* cv. Mitchell.**

**Supplemental Figure S4. Effect of transient suppression of *EVER (AETF5)* in petunia cv. Mitchell flowers on volatile emission.**

**Supplemental Figure S5. Suppression of *EVER (AETF5)* in petunia line P720 flowers leads to enhanced volatile emission.**

**Supplemental Figure S6. Application of exogenous GA upregulates expression of *EVER*.**

**Supplemental Figure S7. Protein sequence alignment of EVER from different petunia species.**

**Supplemental Figure S8. Phylogenetic tree representing the relationships between R2R3-MYB subgroup 1 members and other R2R3-MYBs.**

**Supplemental Figure S9. Protein sequence alignment of EVER from Petunia cv. Mitchell with other subgroup 1 members.**

**Supplemental Figure S10. Generation of *Cas9*-expressing petunia plants with optimal nuclease activity.**

**Supplemental Figure S11. Evaluation of the molecular nature of *Cas9*-transgenic regenerated petunia plantlets following inoculation with pTRV2-sgRNA-*EVER*.**

**Supplemental Figure S12. Generation of *Cas9*-expressing pepper plants.**

**Supplemental Figure S13. Headspace, internal pools and GA treatment of *ever* knockout lines.**

**Supplemental Figure S14. Enriched GO “Biological process” terms in *ever* knockout line #2.**

**Supplemental Figure S15. Enriched GO “Biological process” terms in petunia petals transiently overexpressing *EVER*.**

**Supplemental Figure S16. Knockout of *EVER* alters fatty-acid content in wax of petunia ‘Mitchell’ petals.**

**Supplemental Figure S17. *EVER* knockout does not affect cell morphology or cuticle permeability.**

**Supplemental Figure S18. *EVER* knockout does not affect cuticle thickness.**

**Supplemental Figure S19. Knockout of *EVER* does not affect epicuticular wax load in leaves.**

**Supplemental data and tables**

